# Hydroplane I: one-shot probabilistic evolutionary analysis for scalable organizational identification

**DOI:** 10.1101/2025.10.14.682387

**Authors:** Connah G. M. Johnson, David D. Pollock

## Abstract

Applying evolutionary genomics to microbial and viral community sequence information presents significant challenges. Metagenomic sequences are typically stored in large databases of short-read fragments with unknown relationships. Available tools for analysis are often slow, rely on incomplete subsets of data, or focus on narrowly defined sub-problems. Moreover, existing tools often depend on simplistic model assumptions or treat inferences as empirical data, which can distort downstream analyses. In this work, we present theory and initial validation of the first stage of a fast, one-shot Bayesian approach to metagenomic evolutionary analysis. Our primary result here demonstrates effective use of simple models in experimental design using collections of short proximal short oligonucleotide sequences (*kmers*) to detect probable homologs, and validates the speed, sensitivity and specificity of this approach in a test set of whole *Prochlorococcus* genome sequences.

**Significance Statement:** The Hydroplane algorithm provides an efficient framework for analyzing co-evolutionary relationships among microbial and viral genomes using metagenomic data. Hydroplane guides the identification of homologous regions, estimates lineage diversity, and reveals evolutionary events such as mutation, divergence, selection, and horizontal gene transfer. Its computationally lightweight design supports scalable genome clustering and adaptation analyses, enabling the study of rare microbial and viral genomes across diverse ecological contexts. With an accessible implementation, Hydroplane broadens the scope of evolutionary genomics research, offering critical insights into host-pathogen dynamics and supporting biopreparedness through the study of microbial and viral genetic relationships.

## Introduction

Understanding how microbial and viral lineages diversify and adapt to different environments requires detailed modeling of the evolutionary dynamics of individual genes, particularly to detect convergence and coevolution. This effort depends on a comprehensive understanding of lineage diversity, including rare organisms that may not frequently occur in a given environment. Recent studies have evaluated the performance of algorithm implementations in codified metagenomics tasks such as assembly, genome and taxon recovery through binning, and taxonomic profiling, with a focus on identifying rare genomes [1]. Although the catalogued tools provide numerous options, as noted by Koslicki and colleagues, few offer robust methods for uncertainty estimation in their results, and even fewer provide clear mathematical models for the theory underlying their approaches [2]. The YACHT algorithm, developed by Koslicki and colleagues, stands out for its clear mathematical foundation, computational efficiency, and strong performance[2].

As demonstrated in *YACHT*, alignment-free *kmer* approaches are crucial for creating scalable metagenomics analyses, and a sound theoretical framework is essential. However, efficient methods exist for rapid analysis of repeat structure in mammalian genomes [3], [4], [5], [6], [7], which are on the order of 1000x larger than microbial genomes, and 100,000x larger than many viral genomes. These methods are optimized to handle large-scale data effectively, and seem suitable for adaptation to studying diverse microbial and viral lineages.

Here we describe Hydroplane’s overall goals, a software development project designed to analyze homologous regions among closely related bacterial and viral lineages using well-developed statistical methodologies in a multiple comparison framework. As with our earlier Eukaryote repeat analysis, the concept centers on clustering observed *kmers* into groups called *pclouds*, which are highly similar and repeatedly observed in homologous genome segments. We detect homologous regions by evaluating the spatial proximity, along a genome or genome fragment, of *kmers* belonging to *pclouds* that are known to be adjacent in already acquired target regions.

In the current manuscript (Hydroplane I), we focus on how homolog detectability depends on the balance of *kmer* length, homolog divergence, target segment length, target representation depth, and the related size of the set of *pclouds* used to identify that target. We validate that mathematical predictions using a simple evolutionary model provide useful approximations for experimental design applied to real genomic data, a test set of *Prochlorococcus* genomes. For these genomes, we provide observed genome-wide detection success for short-read sized fragments, demonstrate rapid visualization of inconsistency in divergence rates along these genomes, and discuss implications for future *pcloud* design and detection in metagenomic sequences.

Special attention was given to ensure all genome-wide evaluations and many preliminary calculations were performed with *O*(*L*)complexity, where L ∈ N represents the total length of all sequence reads in a sample set. This optimization allows us to handle moderately large datasets on standard desktop or laptop computers, making the analysis accessible to most users. Memory usage is also structured efficiently through tunable *kmer* memorization. Hydroplane algorithm is implemented in the highly readable *Go* programming language, which facilitates easy comprehension of the algorithmic structure, minimizes reliance on complex hierarchical libraries, and provides a straightforward API and file-based command structure. Compiled versions of Hydroplane are available for MacOS, Unix, and Windows systems, further improving accessibility, replicability, and usability. These features collectively support scientific advancement in a rigorous model-based approach to evolutionary metagenomics that avoids treating inferences as empirical data.

## Results

### Detection and divergence

Detecting homologous sequences in metagenomic samples depends on the amount of divergence between the homologous sequences and available reference sequences. Divergence also affects whether detected sequences are accurately placed in a phylogenetic or classification context. Similarity, the opposite of divergence, is often quantified using the average nucleotide identity (*ANI* ∈ *R*) among genes. In this case, divergence, *δ*, might be estimated from ANI as 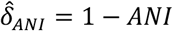. However, divergence rates vary along genomes, and also vary among genomes in metagenomic samples, such that static measure like ANI or 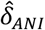 may not be as useful as parametric distributions such as e.g. 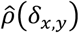, the estimated distribution of divergence among sites in the genome between two organisms *x* and *y*, or the estimated distribution of distances among all members of two groups, 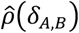.

#### Clouds, storms, and tempests

Following Koslicki [2], the length of *kmers* used in any part of the analysis will be *k* ∈ *N* nucleotides, the set of all *kmers* of length *k* in a reference genome *i* is *G*_*i,k*_, and the count of each *kmer* in *G*_*i,k*_will be *K*_*i,k*_. This last set is output by *Hydroplane* (labelled as a “*kcount*” file). Similarly, the set of *kmers* across *N*_*_ genomes is 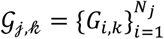. The complete set of locations of these *kmers* in the reference genomes is 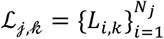, where *L*_*i,k*_ is the set of locations for *kmers* of length *k* in reference genome *I*. This set is optionally output by *Hydroplane* as a “*pcounts*” file. To distinguish between full reference genome *kmer* sets and those that have been filtered or “sketched” to produce subsets, we annotate the set of *kmers* in filtered sets with an asterisk, eg,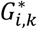.

In previous *pcloud*-based analyses of transposable elements, we introduced the concept of element-specific *pcloud* (*ESP-cloud*) sets such as *Alu-clouds* or *MIR-clouds* that were built from the human transposable elements *Alu* and *MIR* to detect fragments in the human genome that were derived from those transposable elements [3]. In the analyses presented here, we generalize *ESP-cloud* concept to arbitrary-length genomic regions, and the set of clouds for such a region will be called a *stormcloud*, or *storm* for short (Figure 1). Maintaining the weather analogy, a set of *stormclouds* for a longer genomic region, chromosome, or complete genome will be called a *tempest*.

**Figure 1.**
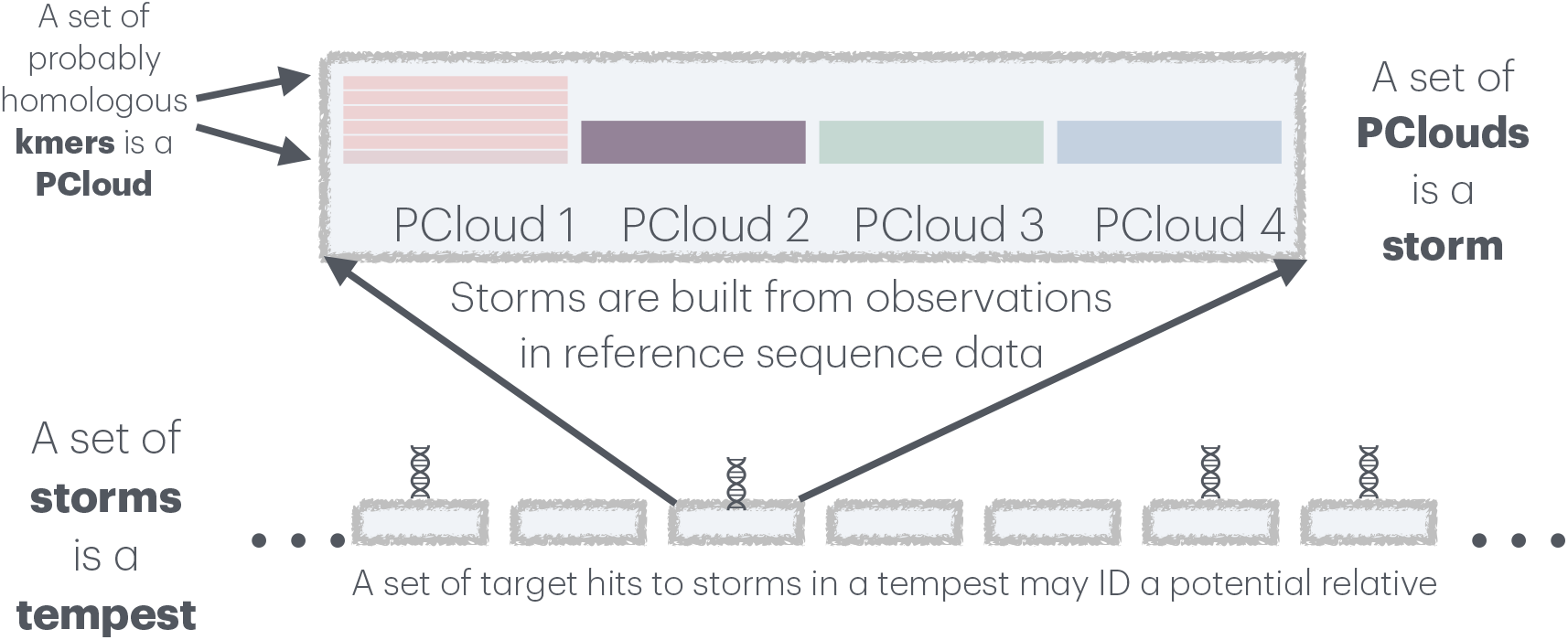
Structure of clouds and storms. The graphic illustrates building up pclouds of observed kmers that are probably homologous based on alignment-free statistical methods. The four clouds illustrated are adjacent but non-overlapping; non-overlap is not a requirement, and the degree of overlap is a user-determined variable, but overlapping clouds should be accounted for in analysis. The storm shown to form a tempest are also non-overlapping, but as with the clouds this is not a requirement, but rather an analytical convenience.

For example, in many analyses presented here, we sketch initial storms from a complete *Prochlorococcus* genome by taking each of the first 20 non-overlapping *12mers* as seeds for their own *pcloud* and clustering these seeds into a *storm*, then proceeding similarly through the remainder of the genome (Figure 1). Applied to a 1.2 Mbp genome, this would result in 5,000 storms, with each storm set to detect non-overlapping fragments of length 240 bp. Note that it is not necessary for the *clouds* or *storms* to be non-overlapping (for example, clouds overlapped by all but 1 bp in the ESP-cloud analyses [4]), but here we use non-overlapping *clouds* and *storms* to ensure independence among them and to ease statistical analyses. Similarly, although *Hydroplane* is designed to build up the *pclouds* and *storms* across multiple genomes, we focus here on the detection properties of *storms* built from single genomes to maintain statistical clarity. We will refer to this sketched set of *kmers* 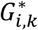 as 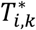 to highlight their organization into *N*_*s*_ storms, each referred to as 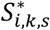 (or just 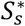 if the *kmer* length and genome context is clear). The creation of *K*_*i,k*_, *L*_*i,k*_, and 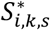 is performed in a single pass through a genome, as is the process of checking a target or query genome for matches to the storms.

### Detection noise

Using individual *kmers* to detect likely homologous sequences can be problematic because short *kmers* may randomly match throughout a genome wherever the accumulation of mutations in non-homologous regions produces identical *kmers* by chance. We can use Koslicki’s and Gu’s simplified model of uniform divergence (UD, [4]) to calculate that the expected number of *kmers* identical to an individual *kmer* of length *k* = 10 by chance is approximately 1.91 in a genome of 2 Mbp, while the expected number drops substantially to only 0.12 for *k* = 12. However, across an entire genome of comparisons, there might then be over 238,000 false positive matches, which is unacceptably large; it is even worse for scans of metagenome samples, which can easily contain over 1 Gbp of sequence. This is the rationale for Koslicki’s approach, which recommends much longer *kmers*, such as *k* = 31 or *k* = 51. Uneven nucleotide and dinucleotide frequencies and other effects can impact the details of this simple calculation, but we observe in *Prochlorococcus* genome comparisons that they are roughly predictive (Figure 2).

**Figure 2.**
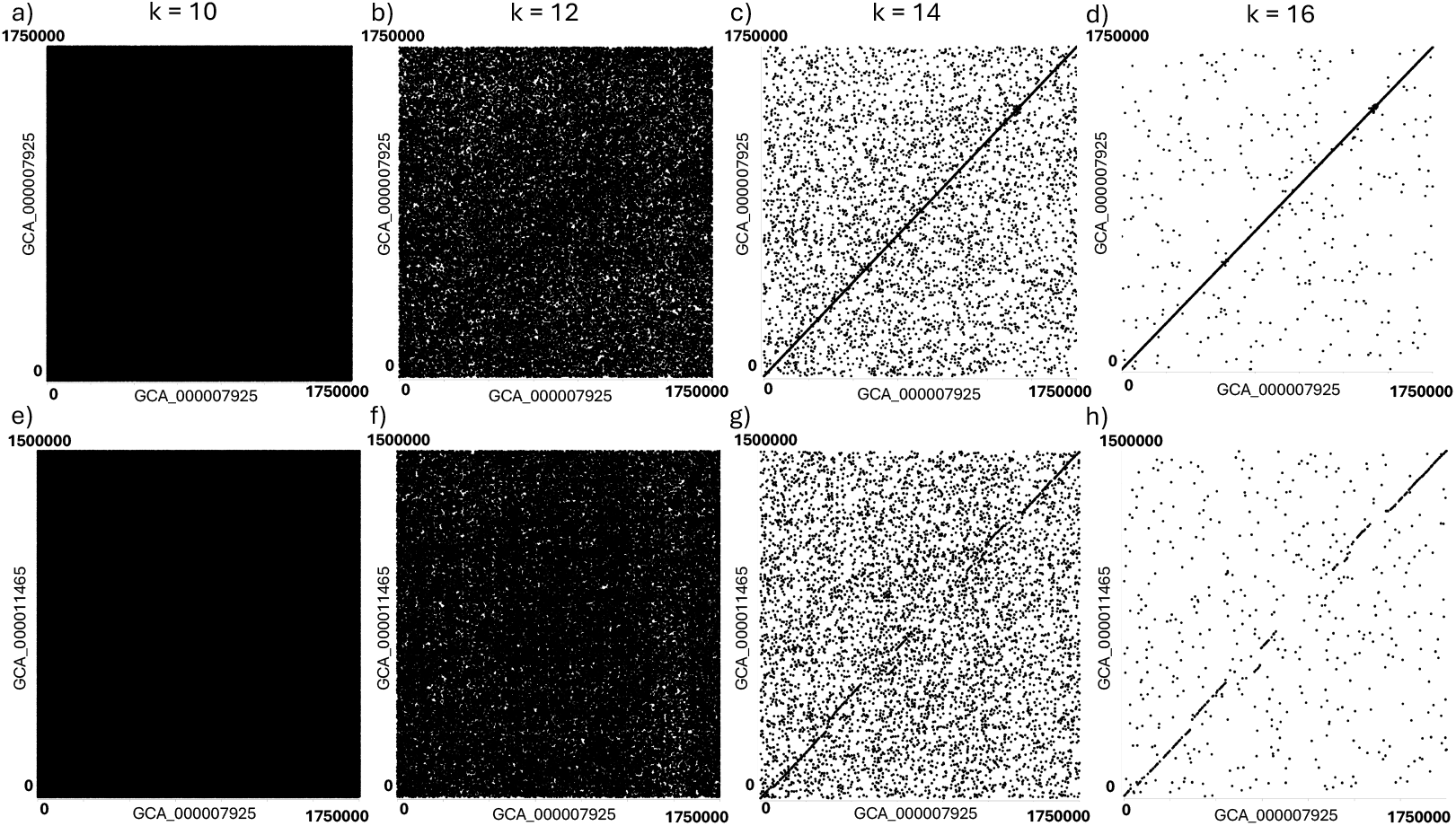
Single *kmer* matching between genomes. The matching between single *kmers* between genomes is shown for 10, 12, 14, and 16 bp *kmers* (a and 3, b and f, c and g, d and h, respectively) for Prochlorococcus genome GCA_000007925 versus GCA_000007925 self-comparison (a-d) and GCA_000007925 versus GCA_000011465 (e to f). Note that off diagonals may occur due to gene or local duplications, or repeat elements, as well as mutational noise. In Hydroplane, the single *kmer* matching was performed through setting the *storm size* to 1, *extendtoside* to 0, and *goodmin* to 1.

For an example storm of size 20, the amount of noise (number of random non-homologous matches) is 20 times worse than for a single *kmer*. This sounds bad at first, but the key to using storms for homology detection is to require more than one storm match within a short region. The chance of observing a second cloud *kmer* by chance in the same 240 bp region as a first random match is extremely small, offsetting the expected ∼4.77 M random *12mer* matches in a 2 Mbp genome, such that only 68.2 double matches are expected. If that is too many, increasing the requirement to 3 matches within a 240 bp region decreases the *genome-wide expectation* of random matches to 0.02. The specificity is very good and the expected false discovery rate is small. Alternatively, increasing the *kmer* length to *k* = 14 has a genome-wide expectation of 0.27 random double matches. Again, we can observe in *Prochlorococcus* genome comparisons that such cutoffs are roughly predictive in noise reduction (Figure 3).

**Figure 3.**
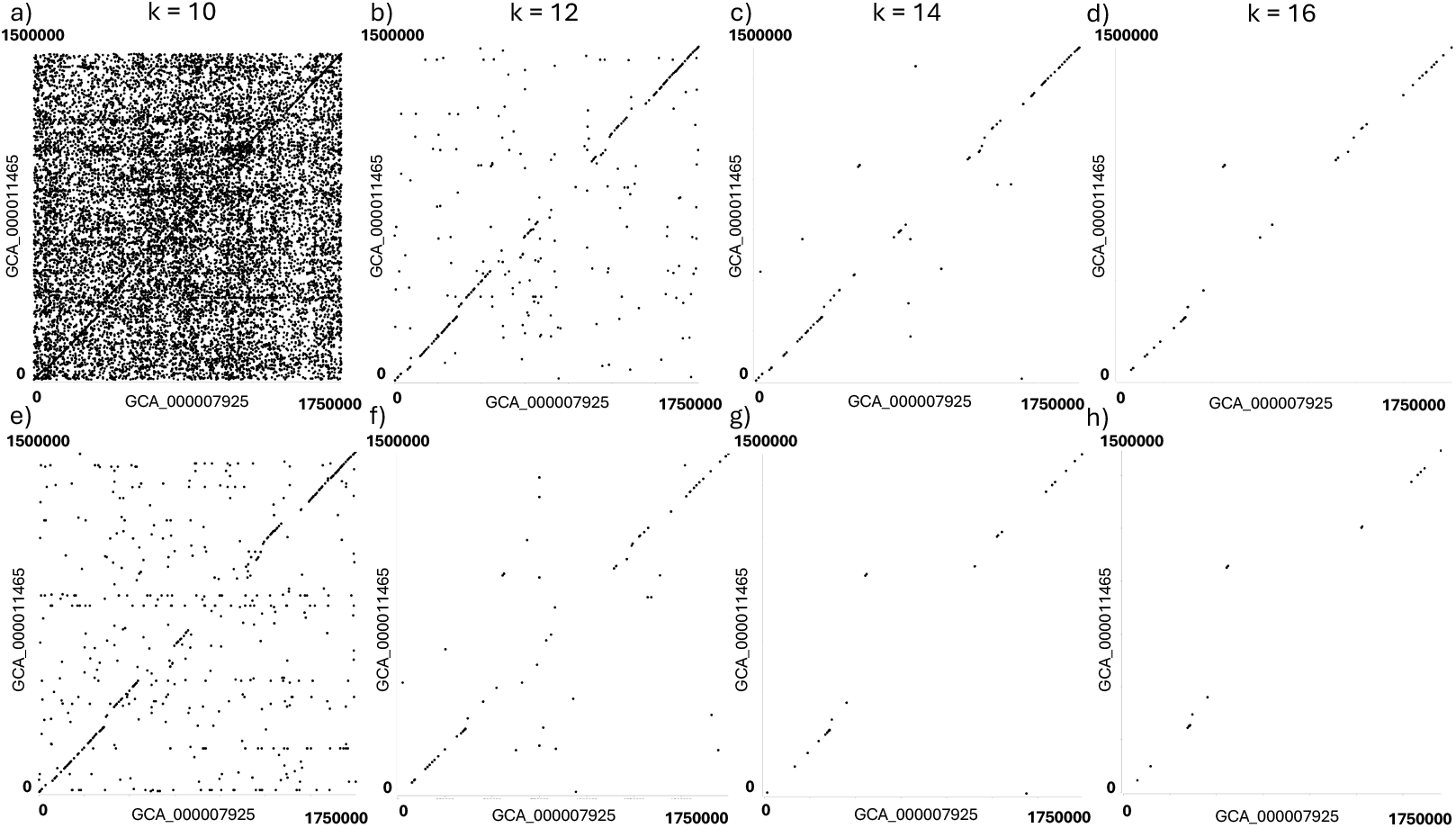
Multiple *kmer* in storm matching between genomes. The matching between multiple *kmers* between the Prochlorococcus genomes GCA_000007925 versus GCA_000011465 is shown for 10, 12, 14, and 16 bp *kmers* (a and e, b and f, c and g, d and h, respectively) with a *goodmin* of 2 (a-d) and 3 (e to f). Note that off diagonals may occur due to gene or local duplications, or repeat elements, as well as mutational noise. In Hydroplane, these multiple *kmer* matchings were performed through setting the *storm size* to 20, *extendtoside* to 10, and *goodmin* to 2 (for the top row, a-d) or 3 (for the bottom row, e-h).

### Signal degradation with time

Differences between homologous genomic regions are expected to accumulate over time due to broad-sense mutation, genetic drift, and selection. While many factors (including degree of neutrality and conservation related to function) affect this process at the level of individual sites and genes, we can use the overly-simplified UD model [2, 4] to calculate expected effects on *pcloud* matching and *storm* hit counts applied over a region, fragment, or *kmer*-length oligonucleotide. If the uniform divergence rate per position is *κ*, then the probability that a *kmer* of length *k* will not have diverged is (1 − *κ*)^*k*^. Because this is the probability that a *kmer* of this length will be detected based on perfect matching, this is the *kmer*-specific sensitivity or recall rate of the procedure.

The sensitivity to detect single *kmers* under UD strongly depends on both *κ* and *k* (Table 1 and Supplementary Data 1). A *kmer* length of 12, for example, may be useful to detect divergence rates as high as 0.2-0.3, with respective detection probabilities for one copy *P*(*d* = 0, *c* = 1|*k* = 12) from 6.9% down to 1.4%, and should be highly effective at divergence rates of 0.1 (28.2% sensitivity) or less. In contrast, longer *i* lengths such as *k* = 24 (as is default in the program *non pareil* [8]), or *k* = 31 and *k* = 51 as recommended by Koslicki, are only highly effect at divergence rates as small as 0.02. Thus, the high specificity of long *kmer* lengths is balanced by poor sensitivity at even moderate levels of divergence. We can infer the effective divergence rate for a homologous region *r* under this model, 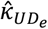, from observed *kmer* detection rates, *P*_*k,r*_, as 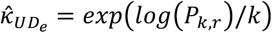.

**Table 1.** Sensitivity to detect *kmers* under the UD model.

| $\kappa$ | $1 - \kappa$ | $k$ | | | | | |
| --- | --- | --- | --- | --- | --- | --- | --- |
|  |  | 10 | 12 | 14 | 24 | 31 | 51 |
| 0.02 | 0.98 | 0.81707 | 0.78472 | 0.75364 | 0.61578 | 0.535 | 0.357 |
| 0.1 | 0.9 | 0.34868 | 0.28243 | 0.22877 | 0.07977 | 0.038 | 0.005 |
| 0.2 | 0.8 | 0.10737 | 0.06872 | 0.04398 | 0.00472 | 0.001 | 0.000 |
| 0.3 | 0.7 | 0.02825 | 0.01384 | 0.00678 | 0.00019 | 0.000 | 0.000 |

We can also consider a conserved sites model with uniform divergence rates at variable sites (*CUD*). For example, if we assume that 2/3 of nucleotides are conserved (*CUD*_4_), which might better approximate a conserved protein region, the results are similar to the *UD* model but with slower dropoff sensitivity with higher mutation rates (Table 2, Supplementary Data 1). Nearly one quarter of 12mers would be identified with a divergence rate of 0.3, for example. We can infer the effective divergence rate for a homologous region *r* under this model, 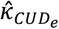, from observed *kmer* detection rates, *P*_*k,r*_, as 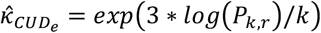, where two out of three sites have rate 0 and one out of three sites have rate 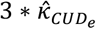.

**Table 2.** Sensitivity to detect kmers under the CUD model. Note that *κ* is the same as in Table 1, *κ*_CUD_ = *κ*_UD_, but because only 1/3 of sites are variable, the average divergence across all sites including conserved sites would be 1/3 as much as the UD model.

| $\kappa$ | $1 - \kappa$ | $k$ | | | | | |
| --- | --- | --- | --- | --- | --- | --- | --- |
|  |  | 10 | 12 | 14 | 24 | 31 | 51 |
| 0.02 | 0.98 | 0.935 | 0.922 | 0.910 | 0.851 | 0.812 | 0.709 |
| 0.1 | 0.9 | 0.704 | 0.656 | 0.612 | 0.430 | 0.337 | 0.167 |
| 0.2 | 0.8 | 0.475 | 0.410 | 0.353 | 0.168 | 0.100 | 0.023 |
| 0.3 | 0.7 | 0.305 | 0.240 | 0.189 | 0.058 | 0.025 | 0.002 |

### Variation in divergence rates in *Prochlorococcus*

To evaluate *Hydroplane* methodology and assess variation along the genomes, we downloaded 109 *Prochlorococcus* genomes from NCBI on Nov 18, 2023 (Supplementary Data 2). Running a single genome against all the others using *k* = 12 and storms with 20 *pclouds* (storm building on ∼2 Mbp, comparative analysis of ∼220 Mbp) took approximately 7 minutes on a 2024 MacBook Air M3 with 16 GB of memory, demonstrating that the algorithm is speedy enough to be useful for local analysis. The complete pairwise run with 109^2^ genome comparisons took 9 hours with *storm size* of 1000, *boundary* of 6000, *extendtoside* of 500, and *goodmin* of 5.

We created distance matrices for all comparisons of both the proportion of storms with hits and the average number of *pcloud* matches among storms with hits (with a hit defined as matches above a preset match minimum (referred to as *goodmin*), 3 unless otherwise specified). *Prochlorococcus* is unusual among genus-level taxon groups in that it is common to observe members of the genus sharing less than 70% similarity [9]. Because of this, low levels of *kmer* sharing were expected with this preliminary hydroplane version (without expanded *pclouds*), and this is what was observed, with many comparisons having proportions of storm hits as low as 0.001 (Supplementary Data 3). To visualize this, we focused on the 27 genomes that contained a single subsequence and increased the accuracy of storm hit frequencies by running storm size 200 and *boundary* = 1200, but still identifying homologs based on only the focal *pcloud* and the 10 *pclouds* on either side. From these results, we created neighbor-joining (NJ) trees (Fig 4), in which the saturation of the *kmer* matching metrics is observable from the poor resolution of deep branches. However, resolution is good within individual clusters, which roughly match clusters observed in previous analyses based on concatenation of gene alignments (Supplementary Figure 1; [9]).

**Figure 4.**
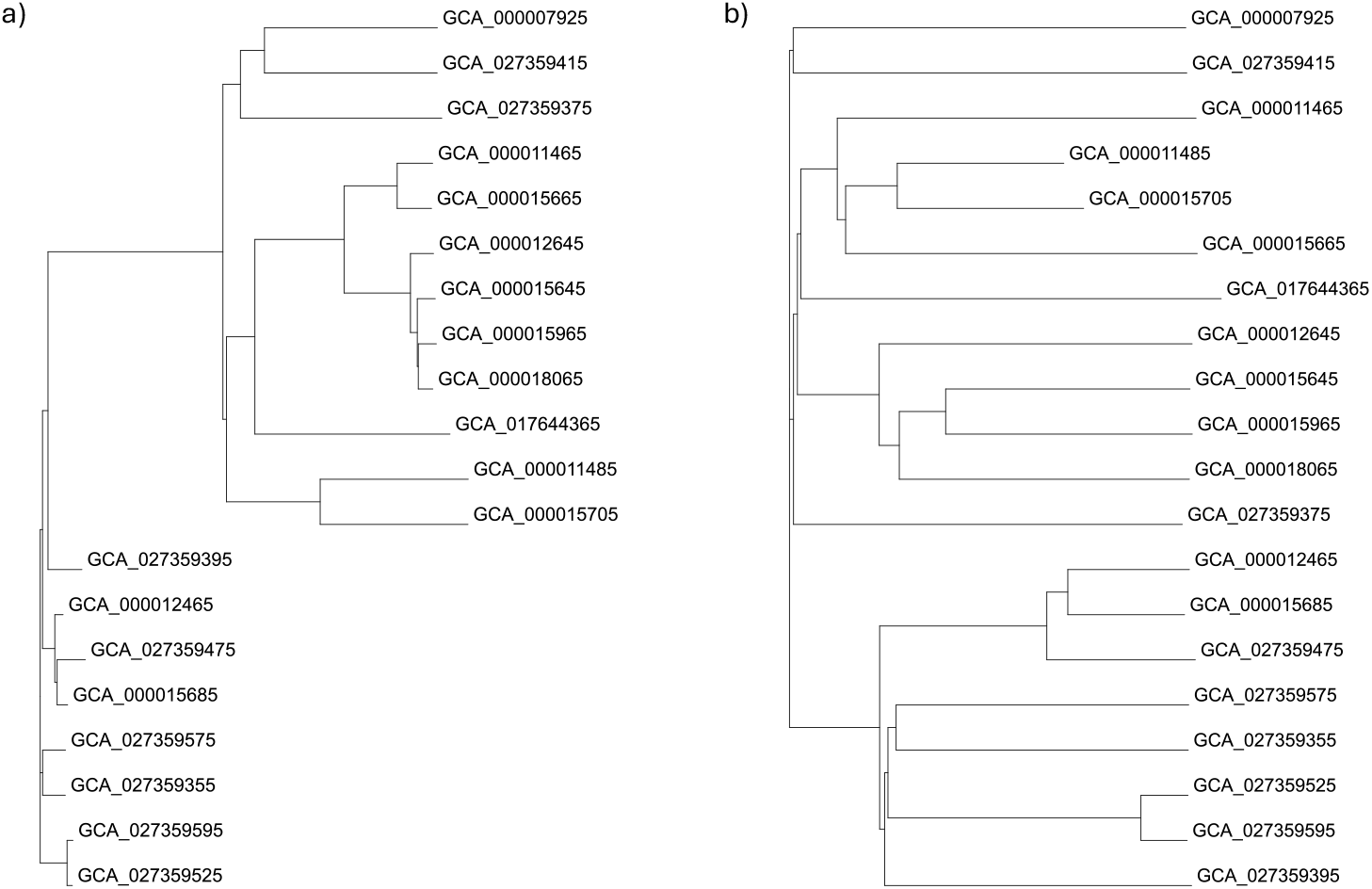
Phylogenetic trees based on proportion storm hits and average matches among hit storms. Distance matrices (Supplementary Data 3) were used to create NJ trees. a) tree based on proportion of storm hits; b) tree based on average matches among storms with good hits.

**Figure 5.**
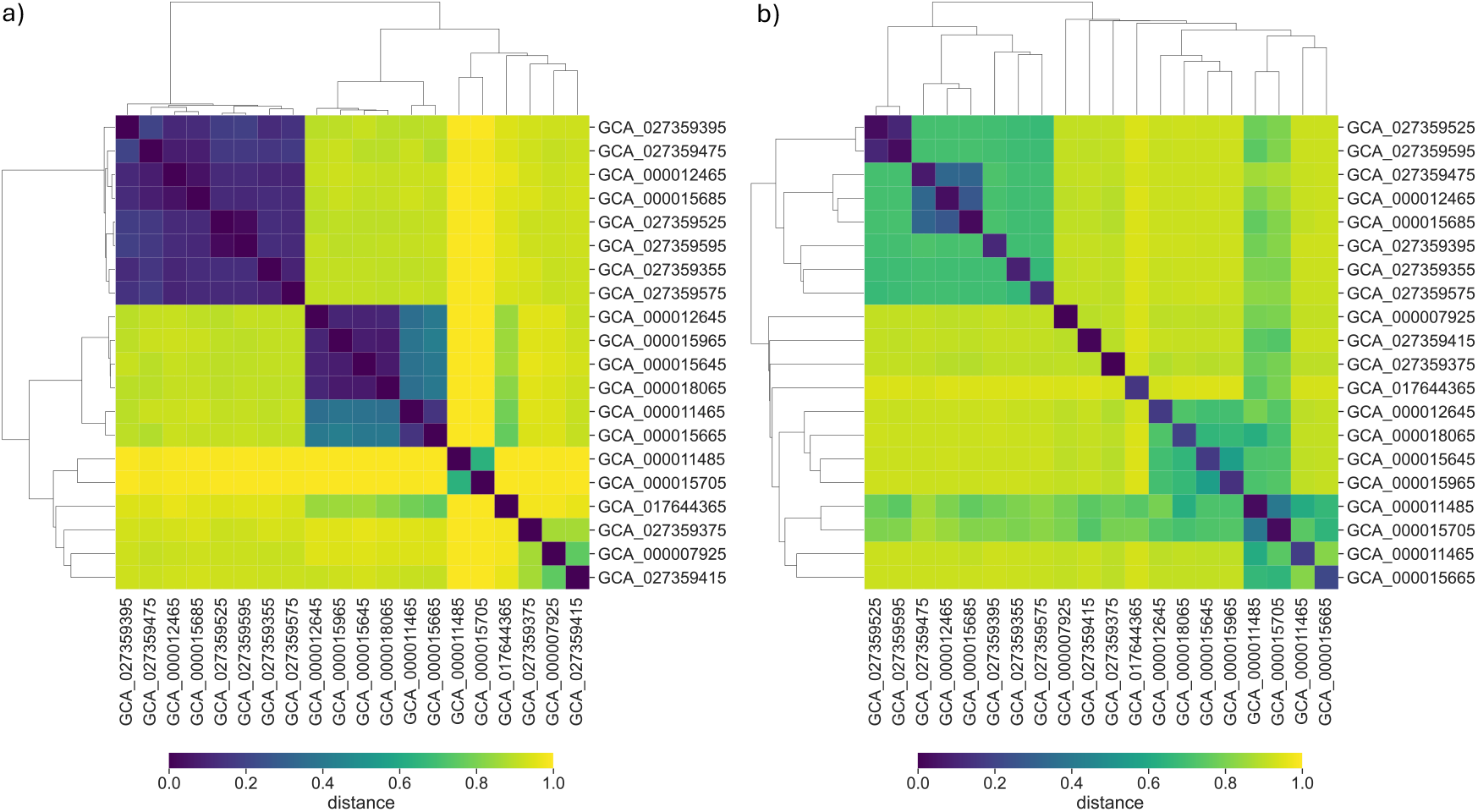
Divergence heatmaps showing distances between genomes. Heatmaps were produced using the distance matrices (Supplementary Data 3) a) heatmap based on proportion of storm hits; b) heatmap based on average matches among storms with good hits.

We note that among storms with hits, the average number of matches in any comparison was often considerably higher than expected based on the proportion of storms hit. This was true for highly divergent genome comparisons as well as more moderately divergent genomes. This indicates that the probability of a match correlates with the probability of matches in surrounding regions. In other words, there is regional variation in divergence rates along the genome. This variation is illustrated by consistent divergence patterns among moderately divergent genomes (Figure 6b), as well as among highly diverged genomes (Figure 6c).

**Figure 6.**
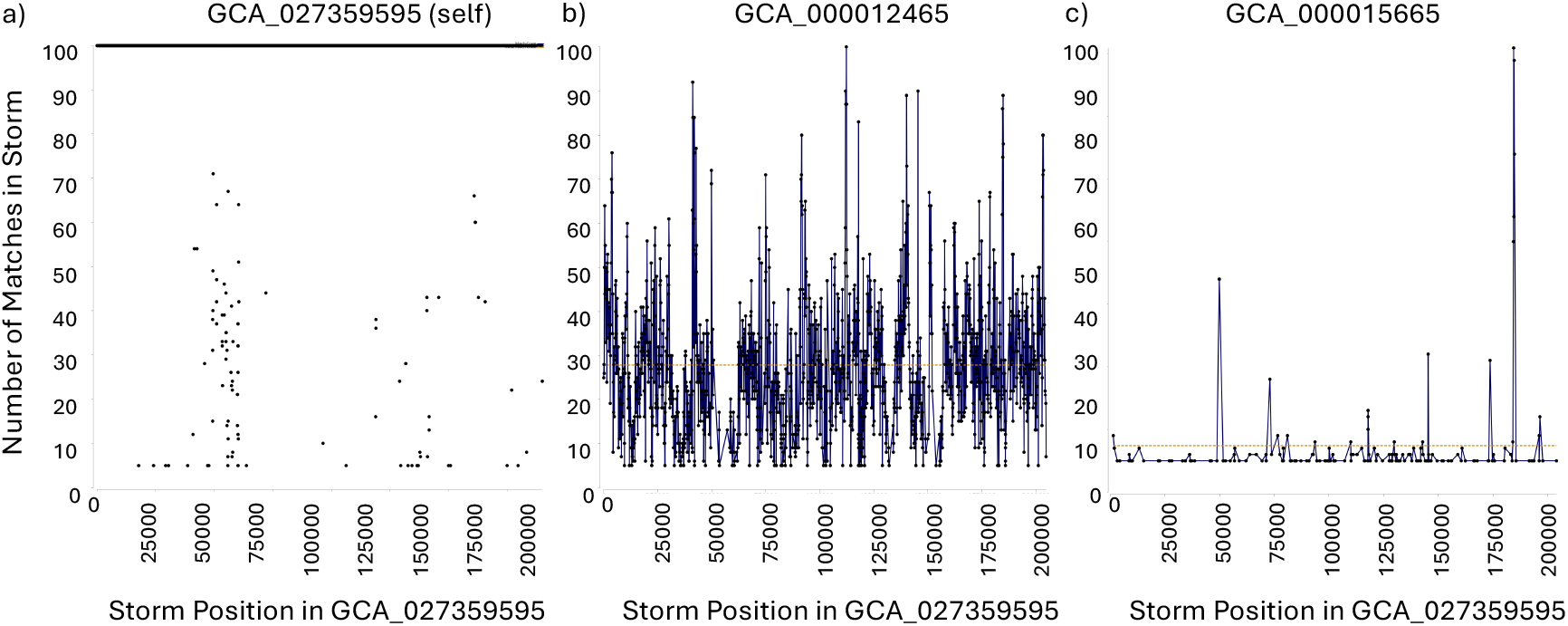
Divergence patterns among highly and moderately diverged genomes. Based on the trees in Figure 4, divergent genome pairs were chosen to illustrate divergence patterns of cloud sharing. The number of matches in each storm demonstrate the quality of storm identification. These figures are produced with *k* = 12, *storm size of 100, boundary* of *6000, extendtoside* of 10, and *goodmin* of 5. a) The self comparison showing a remapping of storms from the reference to itself. The high storm matches equal to the storm size show complete mapping as expected. The lower storms suggest the presence of repeats or sequence noise. b) Patterns of divergence within moderately diverged genome clusters. Fewer storms are seen to be completely mapped with divergence across these storms and regions of low storm mapping. c) The pairs were chosen to have diverged near the base of the trees in Figure 4, with no shared branches among them. Storm maps are fewer and of low quality. However, spikes of high quality storms are identified suggesting strongly conserved regions.

### Deep Analysis of a pairwise genome comparison

To more closely examine detection rates, estimate rates of divergence, and validate homology detection, we chose in-depth comparisons of GCA_027359595 (ASM2735959v1, Prochlorococcus marinus str. MIT 0912) to build storms and GCA_000012465 (ASM1246v1, Prochlorococcus marinus str. NATL2A) as a query sequence to detect homology to storms. From Figure 6b, the mean *pcloud* match rate was 28.69% across the resulting ∼1600 storms (Figure 6). This corresponds to 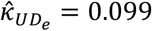 and 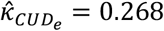, or ∼10 − 27% divergence at variable sites. However, the match identification rates varied from 10-97% along the genome, with some of the highest rates corresponding to the rRNA-coding regions (Figure 6).

The variance of match frequencies is much higher than expected for a binomial with *N* = 100.

## Discussion

Here we have successfully validated that, using *Hydroplane* methodology, we can in theory and practice rapidly detect a high proportion of likely homologous short 240 bp bacterial sequences in the range of 0.2-0.3 *κ* per site divergence from reference sequences, using *Prochlorococcus* reference genomes for demonstration. This is well beyond the expected range of divergence for most bacterial groups (although in picocyanobacteria there are many pairwise comparisons deeper than this [9]), demonstrating its utility to collect sequences involved across a range of moderately recent evolutionary processes. Our theoretical analysis makes it clear how and why one can obtain high sensitivity and specificity with short (10-14 bp) rather than long (e.g., 24, 31, 51 bp) *kmers*, while retaining speed in the absence of alignment or phylogenetic reconstruction.

Although there is considerable frequency variation in mono- and dinucleotides, we found that expectations of noise based on simple models of random sequence generation were useful for parameter control. Likewise, although there is considerable variation among sites and regions in the amount of divergence throughout genome comparisons, expectations of signal based on simple models of sequence divergence per site were useful to understand the variation among sites and between genomes with varying divergence levels. One way of putting this is that the mathematical result of multiplying small numbers (here, the probability of randomly obtaining multiple copies of a small subset of possible *kmers* within a short stretch of sequence) is an exceptionally small number. This multiplication effect dominates the accumulation of randomly misidentified *kmers* that depends linearly on sequence length.

This manuscript is focused on the first stage of a fast, one-shot Bayesian approach to metagenomic evolutionary analysis, and the next stage will focus on both the theory and practice of building up the *<u>pclouds</u>* for homology detection. In conjunction with this, we will analyze the iterative process of adding newly identified and localized *kmers* into clouds for enhanced detection of homologs within and across metagenomic samples. This second stage will clarify and practicality of controlling the goal of purely computationally linear alignment-free comparison of large numbers of sequences, as well as the practical benefits of utilizing clouds of observed likely homologous *kmers* to identify homologous segments in bacterial sequences.

Because the purpose of this initial stage is focused on the fundamental mathematics of quick probable detection of homologs, we note that it is designed to be applied flexibly according to user goals and preferences. The data presented is aimed at fairly high signal to noise ratios (that is, few false positive predicted homologs) when applied to 2 Mbp genomes, but in many cases such signal to noise ratios may be excessive. If subsequent analyses are designed to allow for uncertainty, for example, it is reasonable to allow much higher levels of false positives. If one is interested in asking questions such as “how many of the reads in this metagenomic sample match this genome?”, you want to limit the number of false negatives as well. With the estimates of false positive rates and false negative rates (depending on site-specific divergence rates and patterns) that this methodology can deliver, one can obtain reasonably unbiased estimates of such fractions (see de Koning et al. for example [4]). Unlike vertebrate genomes, which are dominated by variably divergent repeats and duplicated genes and regions, bacterial genomes (at least *Prochlorococcus*) appear to have relatively stable and empirically estimable false positive rates, which makes such analyses easier. An immediate goal is to obtain rapid estimates of read fractions using *Hydroplane* that account for variation in divergence rates along genomes and variation of realized divergence among organisms in a metagenomic samples.

Another reason to allow moderately high false positive rates in initial *Hydroplane* analyses is that there are many avenues to refine homology estimates using *Hydroplane* with different run parameters and calculations. To start with, if one has two moderately short regions that are thought to be homologous, the Hydroplane approach can in principle be run with *storms* made from overlapping offset-by-one *kmers* to seed the *pclouds*. Overlapping and directly adjacent *pcloud* matches can then be merged to produce sets of contiguous *pcloud*-matching regions of certain lengths separated by gaps of non-*pcloud* matching regions of certain lengths. Following Wade [7], the chance probability of such a combination of matching regions with gaps can be calculated with much higher precision than the fast non-overlapping analysis examined here. These calculations can also include whether the order of the putatively homologous regions is similar to the order of the storms that they match.

A further useful independent source of information is synteny (Ahrens et al 2021 [10]). For example, if a focal region is surrounded by two matching homologous regions, there is a high probability of homology in the focal region for that reason alone. This dramatically changes Hydroplane null probability calculations. For example, given a syntenic 250 bp region, the probability that a particular *kmer* of length 6 occurs by chance is only about 6%, and thus the probability of two independent *kmers* of this length by chance is only 0.37%. Using shorter *kmers* in this context will greatly increase sensitivity without overly increasing noise.

We did not include storm match order analysis here because it slows down analysis, but it is perfectly acceptable as a post-analysis filtering methodology (Wade 2022 [7]) and will be particularly impactful in comparisons that are still uncertain even with moderate numbers of matches (at least 3-5). Such uncertainty with moderate match numbers can happen with sparse matches in long regions or with short *kmers* even in short regions.

Here, we have only briefly touched on differentiating sequences that might code for proteins (i.e. the *κ*_*CUD*_ calculations) or other functional molecules. This is because we expect that 1) protein variation patterns will be naturally accounted for when we build full *pclouds* from diverse data; 2) we are designing the Hydroplane project to derive conclusions based as much as possible on first principles rather than relying on treating past inference (such as what sequences code for proteins) as data; 3) we are focused on detecting homology rapidly in moderately diverged sequences, whereas protein comparisons are more critical for deeper comparisons; and 4) adding proteins would increase program complexity and slow down computation (something we do not want when aiming to address Tera bp or even Peta bp of data).

That said, once full *pclouds* are constructed, future development will likely focus on a few promising avenues. It may be useful to consider adding *pclouds* that mix observed variation within a *pcloud*. For example, if neutral variation at two different 3rd codon positions were observed in separate instances, it might be predictable that double variants are likely but not yet observed. Another option is to build *pclouds* from reference genomes translated amino acids, then translate them to nucleotide sequence *pclouds*. This could be done based on annotations or could be done for all 3 codon position offsets in both directions (see Wade 2022 [7]), again reducing dependency on possibly erroneous or incomplete annotations. Finally, it may be interesting to use *kmers* that are interspersed with one another, offset by 3. Such an approach might reliably identify segments conserved at 1st or 2nd codon positions without a great deal of preliminary calculation. All of these approaches should be tested to evaluate their sensitivity, specificity, and speed for a variety of deeply diverged genomes.

## Materials and Methods

### Hydroplane Program

The *Hydroplane* project is written in the Go programming language, and code, examples, and instructions are available at github.com/NWBRaVE/Hydroplane and github.com/PollockLaboratory/hydroplane. The code is derived from the previously published code base github.com/PollockLaboratory/AnCovMulti. All parameters used in the file can be controlled by three files: factory, mode, and control, which are respectively intended to contain parameter definitions and parameters that are not expected to change, parameters controlling a particular mode of operation, and control parameters that might be regularly changed during testing. In the analyses run for the current manuscript, reference genome input and output are found in the *references* or *output* folders.

After setup, *kmers* of length *k* are read, counted and localized for each file. In the implementation discussed in this paper, *storms* are seeded and collated into a *tempest* based on adjacent non-overlapping *kmers* in the first reference genome read, and the number of matches to each storm is evaluated for all reference genomes. The *storm* size and location of *pcloud* seeds within storms are properties of the *tempest*. Here, seed locations are calculated as half the *storm* size, rounded down to the nearest integer.

Hits between storms and reference sequences are calculated based on identification of a minimum number of identical *kmer* matches to *clouds* of the *storm* in a confined region of the reference sequence (Figure 7). To control the difference between identifying a *storm* match and accurately counting the number of matches to the *storm*, we allow the user to select a subset of *pclouds* at the center of the *storm* for identification. This subset is centered around the seed, and includes a number of clouds on both sides controlled by the *extendtoside* parameter. To speed these comparisons, for query reference genome *j*, locations *L*_*i,k*_ in ℒ_*j*,*k*_, where *k* matches a *k* in 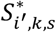 within the identification subset are collected as locations of potential *storm* hits. These potential *storm* hits are identified moving outward to both sides from the seed, with the location of the potential storm given as the location of the first identified *storm kmer* in a region. All *kmers k* in 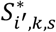, both within and outside of the identification subset, are counted as matches in the potential *storm* hit if *L*_*i,k*_ in ℒ_*j*,*k*_ is within the *boundary* distance (set by the input *boundary* parameter) of the original potential *storm* hit location. Once a storm is parsed, potential *storm* hits are saved as good hits (putative homologs) if the number of identified matches are greater than or equal to the user input *goodmin* parameter.

**Figure 7.**
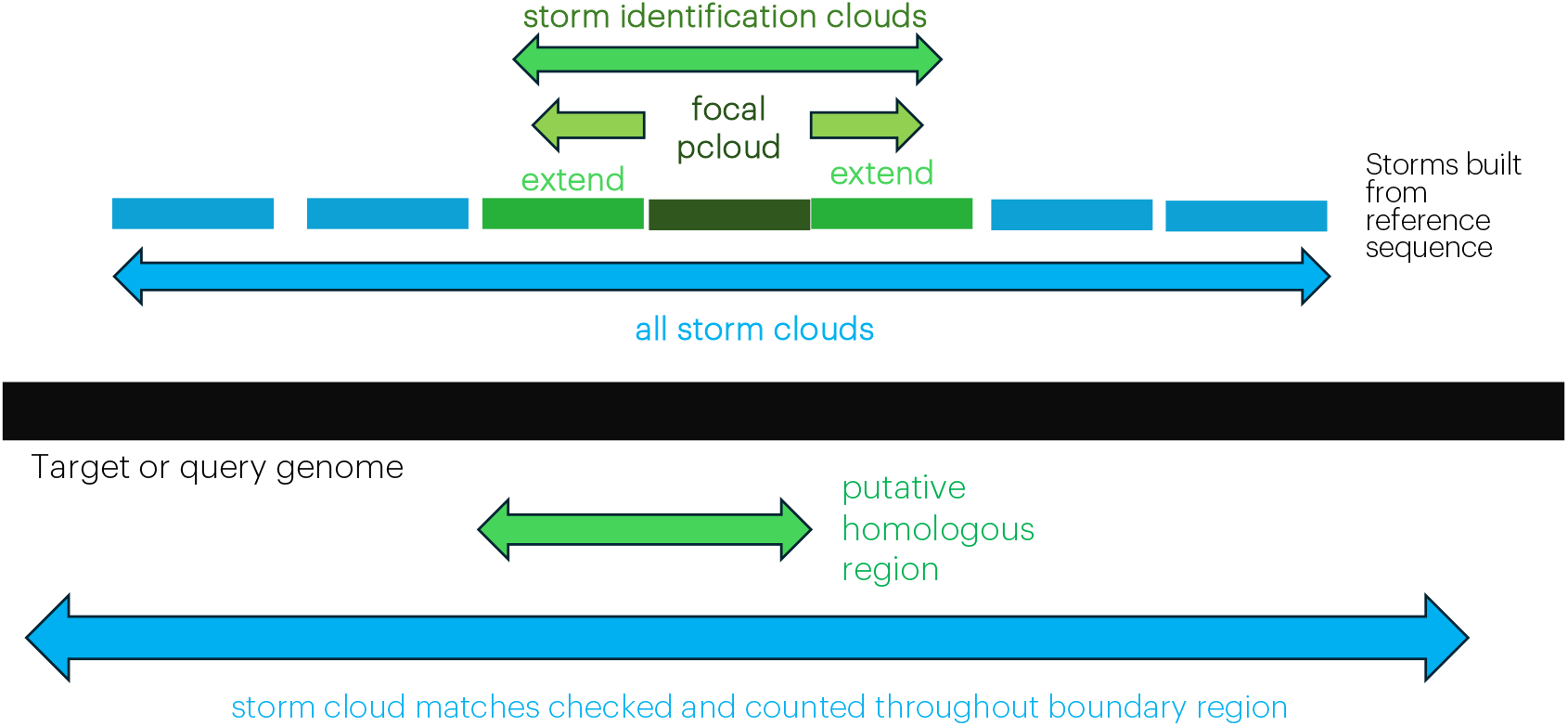
Patterns of homolog identification with storm subsets and validation within boundary. The graphic illustrates how a storm subset is used to identify homologs, allowing validation of homology status within defined boundaries around the focal cloud of the storm. This clarifies the use of the *focal kmer, stormsize, extendtosides*, and *boundary* parameters.

Although this procedure is admittedly slightly complex to explain, it is quite fast. The use of a boundary setting that will scan regions slightly larger than the nucleotide length of the storms allows for insertion and deletion variation between the storm and homologous regions. The use of storm subsets for identification allows us to assess identification rates of the subset while validating identification of homologs and measuring the match rate with longer and therefore more accurate storms.

### Reference genomes

We downloaded 109 complete genomes from NIH/NCBI on Nov 18, 2023 (list in Supplementary Data 2) to use for testing, most of which were approximately 2 Mbp (approximately 2MB). For standardized comparison of two moderately divergent genomes, we used the genome GCA_000007925 (aka ASM792v1, Prochlorococcus marinus subsp. marinus str. CCMP1375) to build storms and GCA_000011465 (aka ASM1146v1, Prochlorococcus marinus subsp. pastoris str. CCMP1986) for comparison as a target or query. We found that 27 of these genomes were composed of a single sequence, and 19 of these were labelled “complete” in the genome label, so graphical comparisons were limited to the single sequence genomes (as our aim here is not to reconstruct genome order).

### Other analyses

Phylogenetic trees based on distance matrices were produced using the neighbor joining algorithm [11] as implemented in the scikit-bio v0.7.0 python package [12]. Trees were visualized with Bio.Phylo [13] of the Biopython package v1.85.0 [14]. Simple graphs and heat maps were visualized with *Go* within the *Hydroplane* program.

## Acknowledgments

The research described in this paper is supported by the NW-BRaVE for Biopreparedness project funded by the U. S. Department of Energy (DOE), Office of Science, Office of Biological and Environmental Research, under FWP 81832. A portion of this research was performed on a project award (Enhancing biopreparedness through a model system to understand the molecular mechanisms that lead to pathogenesis and disease transmission) from the Environmental Molecular Sciences Laboratory, a DOE Office of Science User Facility sponsored by the Biological and Environmental Research program under Contract No. DE-AC05-76RL01830. Pacific Northwest National Laboratory is a multi-program national laboratory operated by Battelle for the DOE under Contract DE-AC05-76RL01830. A portion of this paper was supported by the University of Colorado School of Medicine.

## Data availability

All code and example input and output data are available at github.com/PollockLaboratory/hydroplane1a and github.com/NWBRaVE/Hydroplane

## Author contributions

D.D.P designed the research. C.G.M.J., D.D.P developed the code and performed the analysis. All authors contributed to writing the paper.

